# Obese serum factors aggravate DNA damage, alter DNA damage response, and promote proliferation in colon cancer cells

**DOI:** 10.1101/2025.03.11.642555

**Authors:** Bhavana Deshmukh, Himanshi Yaduvanshi, Firoz Khan Bhati, Manoj Kumar Bhat

## Abstract

**Highlights:**

- Obese serum aggravates DNA damage and reduces levels of DNA repair molecules in colon cancer cells.
- Obese serum activates DDR mechanisms by upregulating P-p53ser15 and pchk2.
- *ATM-/-* mice are prone to AOM/DSS induced colon polyp formation.
- Obese serum contributes to the survival phenotype of colon cancer cells.

Graphical Abstract

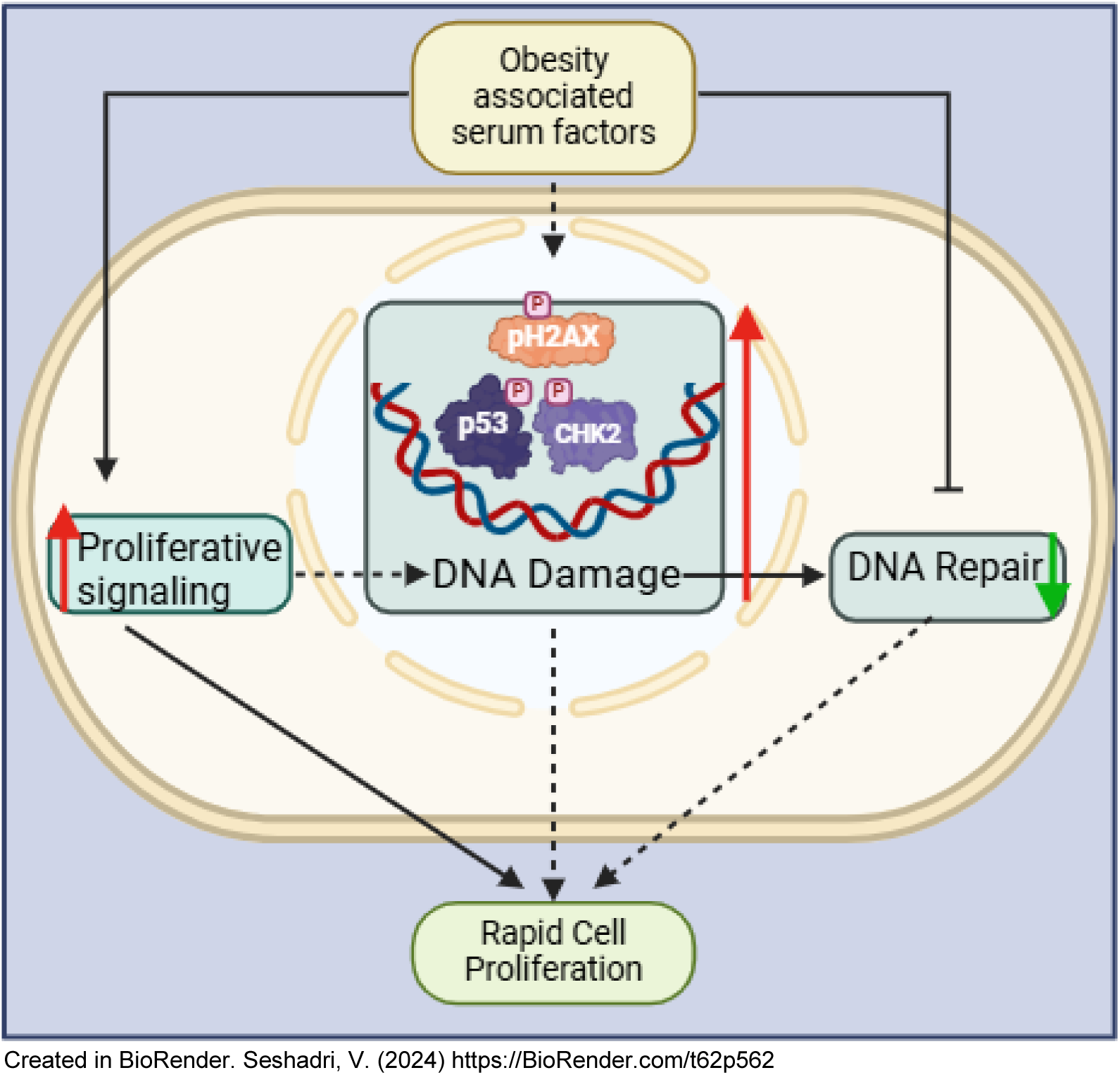

Clinical data indicate a positive link between obesity and DNA damage, which has also been implicated in various pathological conditions. Obesity increases the risk of occurrence and progression of cancers including colon cancer. The underlying mechanisms of the association between obesity-induced alterations in DNA damage response and colon cancer remain unexplored. The present study attempts to investigate the functional status of cellular DNA damage response in an obese environment and its association with colon cancer. To address this, cells were cultured in media supplemented with serum collected from mice fed a normal fat-diet and high-fat diet. Subsequently, the DNA damage response and phenotypic parameters were evaluated. Experimental results revealed that cells cultured in HFD serum exhibited increased DNA damage and reduced levels of DNA repair molecules together with activation of DDR by upregulating levels of pH2AX, P-p53ser15, and pchk2 proteins. Moreover, the cell growth assay revealed a rapid proliferation of cells cultured in HFD serum. Furthermore, HFD mice administered with azoxymethane/dextran sodium sulfate (AOM/DSS) exhibited a higher occurrence of colon polyps compared to normal diet-fed mice. Interestingly, in ATM knockout mice (ATM-a key DDR-related molecule) a higher occurrence of polyps was detected compared to ATM wild-type mice, suggesting a possible association of ATM with polyp formation. Thus, by perturbing DDR, repair pathways, and increasing cell survival, obesity creates a favorable environment for the proliferation of cells. Collectively, this study advances understanding of obesity-altered DDR and its association with cancer cell proliferation.

## 1. Introduction

Cancer and obesity are the modern world’s prominent health concerns, and a significant population is under the grip of these diseases. Higher body mass index (BMI) together with the distribution of fat (central obesity) is considered a major risk factor for cancer. It has been reported that 14% of cancer-related deaths in men and 20% in women are attributed to obesity [1]–[3]. In obesity, excessively accumulated adipose tissue functions as an endocrine organ and secretes many bioactive molecules termed adipokines. Elevated serum levels of adipokines have implications for the pathogenesis of obesity-associated diseases such as hypertension, cardiovascular disease, type 2 diabetes, osteoarthritis, liver diseases, cancer, and kidney failure [4]. Adipokines elicit their effect by binding to cognate receptors, which transmit signals to transcription factors through specific pathways including JAK-STAT, ERK MAP Kinase, etc.

Also, altered serum factors, adipokines, and oxidative stress in obese individuals can create an imbalance in DNA (deoxyribonucleic acid) damage response (DDR) and cell cycle regulation, paving the way for the survival of phenotypically altered or transformed cells. Interestingly, serum 8-hydroxy 2′-deoxy-guanosine (8-OHdG), a marker of oxidative DNA damage, level increases in diabetes associated with obesity, [5]. Also, oxidative stress and DNA damage increase in obese conditions [6]. Moreover, obese individuals often become victims of chronic inflammation. Parameters associated with inflammation are reported to increase oxidative stress in cells and induce DNA damage response [7]. Therefore, the role of obesity-related factors in the pathogenesis of cancer by altering DDR cannot be underestimated.

Obesity increases the risk of developing colorectal cancer by 1.5 to 2 fold [8]. As per the GLOBOCAN 2020 data, colon cancer is the third most prevalent cancer in the world and it also ranked second in cancer-related deaths after lung cancer in both sexes. Estimates indicate that by the year 2035, the mortality from colon cancer will increase by 71.5% [9].

It has been reported that alteration in the DDR cascade fuels genetic instability [10]. When a normal cell is exposed to genetic insult, its DDR mechanism diligently rectifies the faults. Cells having damaged DNA undergo scrutiny at cell cycle checkpoints and are eliminated by apoptosis. If DNA damage is sustained during cell cycle checkpoints, then it can lead to the proliferation of abnormal cells. Many human malignancies also arise due to gene-level mutation in DDR molecules [11].Reports from our laboratory and other groups suggest that obesity accelerates the progression of melanoma [12], [13], and breast cancer [14]. Mechanistically, activation of multiple pathways has been linked to the progression of tumors. Vergoni et al. reported that a high-fat diet (HFD) induces DNA damage, shortens telomere, and causes activation of the p53 pathway in mouse adipocytes [15]. Also, inactivating mutations of DDR-related protein ataxia-telangiectasia (ATM) is associated with a higher risk of breast cancer and a moderate risk of colon cancer [16].

Cancer arises from irreparable DNA damage in normal cells, leading to genetic abnormalities. Genetic lesions and damage activate complex signaling pathways that regulate repair, survival, senescence, and apoptosis. In cancer cells, mutations,oxidative and replicative stress result in increased DNA damage and reduced DNA repair capacity, further driving genomic instability [16]. These cellular fate decisions are intricately linked to cancer initiation and progression. Therefore, investigating the molecular events associated with the DNA damage response (DDR) is crucial in understanding the connection between obesity and cancer. The findings presented in this manuscript enhance our understanding of how obesity-associated factors influence DDR, potentially contributing to cancer development and progression in obese conditions.

## 2. Materials and methods

### 2.1. Cell lines and culture conditions

Murine colon cancer cells, CT26 and MC38 were kindly gifted by Dr. Dipankar Nandi, Indian Institute of Science (IISc), India, and Dr. Ruben Hernandez, Center for Applied Medical Research (CIMA), Spain, respectively. Mouse pre-adipocyte cells 3T3-L1 were obtained from ATCC and maintained in an in-house cell repository at National Centre for Cell Science (NCCS), India. HCT 116 p53 +/+ (Human origin, colon cancer) cells were a generous gift from Dr. Bert Vogelstein, John Hopkins University, USA. MC38, 3T3-L1, and HCT116 p53 +/+ cells were cultured in Dulbecco’s Modified Eagles Medium (DMEM) with 10% fetal bovine serum (Gibco, NY, USA) whereas CT26 cells were cultured in Roswell Park Memorial Institute (RPMI) 1640 medium with 10% newborn calf serum (Gibco, NY, USA). Antibiotics, penicillin (100 U/ml), and streptomycin (100 µg/ml) (Invitrogen Life Technologies, CA, USA) were added in both media. Cells were maintained at 37°C in a 5% CO_2_ humidified incubator (Thermo Fisher Scientific, OH, USA).

### 2.2. Antibodies and chemicals

p53 (sc-6243), phospho-p53ser20 (sc-18079), phospho-ERK (Tyr 204) (sc-7383), ERK (sc-154), survivin (sc-17779) β-actin (sc-1615), β-tubulin (sc-9104) and HRP conjugated secondary antibodies, anti-mouse (sc-2031) and anti-rabbit (sc-2030) were purchased from Santa Cruz Biotechnology (CA, USA). Phospho-H2AX (Ser139) (2577S), phospho-p53ser15 (9284T), and phospho-p53ser15 (9286T) were purchased from Cell Signaling Technology (MA, USA). Phospho-p53ser46 (BD558245) antibody was purchased from BD Bioscience (NJ, USA). Phospho-chk2 (T68) (ab-3501), and Rad51 (ab-133534) antibodies were purchased from Abcam (MA, USA).

### 2.3. Experimental animals and diets

Mice (129S6/SvEvTac-ATM(tm1Awb)/J *ATM+/+* and 129S6/SvEvTac-ATM(tm1Awb)/J *ATM-/-*) were procured from Jackson laboratories (ME, USA). C57BL6/J mice were obtained from the Experimental Animal Facility (EAF), NCCS, India. All mice were inbred and maintained at in-house EAF at NCCS. To induce colon cancer, azoxymethane (AOM)/dextran sulfate sodium salt (DSS) model was developed as per the method described by Singh S. et al., 2020 [17]. Methylene blue staining was performed for confirmation of the dysplastic phenotype of polyp as mentioned by Singh S. et al.,2020 [17]. Obesity in mice was induced by using, a high-fat diet as described earlier [12]. Mice were fed with respective diets for 10-22 weeks until significant changes in body weight and serum profile were observed. The serum was pooled separately from normal diet (ND) and high-fat diet (HFD) fed mice, which was further used for culturing cells. To evaluate the colon cancer incidences in *ATM+/+* and *ATM-/-* mice, both groups were treated with AOM/DSS. After completion of the experiment, mice were euthanized in a CO_2_ chamber, dissected and the colon was excised. Polyps were counted, excised from the colon tissue, and sizes were measured. Tissues were immediately preserved at −80°C until further use.

Diet and water were given to mice *ad libitum*. All the animal experiments carried out were as per the rules and guidelines of the Committee for the Purpose of Control and Supervision of Experiments on Animals (CPCSEA), Government of India. The permission for all animal experiments was obtained from Institutional Animal Ethics Committee (IAEC).

### 2.4. Serum biochemical analysis

Blood was collected from the experimental mice in a collection tube by orbital sinus puncture and centrifuged at 4000 rpm for 10 min at room temperature (RT) to separate serum. Serum triglyceride and cholesterol levels were measured by colorimetric kits as per the manufacturer’s protocol. The kits used to estimate serum triglycerides (MX41032) and cholesterol (TK41021), were purchased from Spinreact (Girona, Spain).

### 2.5. Lysates preparation and immunoblotting

Colon tissues and polyps were excised, chopped, and washed with ice-cold Phosphate Buffered Saline (PBS). Chopped tissues were dis-integrated in radioimmunoprecipitation assay (RIPA) buffer (150 mM NaCl, 20 mM Tris-HCl (pH 7.5), 1 mM Na_2_EDTA, 1 mM EGTA, 1% NP-40 1% sodium deoxycholate, 2.5 mM sodium pyrophosphate, 1 mM β-glycerophosphate, 1 mM Na_3_VO_4_ and 1 µg/ml leupeptin) followed by homogenization and sonication (3 cycles at 21 mA, 10 secs on/off) using sonicator (Vibra-cell, MA, USA).

Cells were scraped from culture dishes and washed with PBS. The suspension was centrifuged at 2000 rpm for 5 min at 4°C to collect cells pellet. Cells were lysed in RIPA buffer. The immuno-blotting was performed as per the protocol described previously by Malvi P. et al., 2015 [13]. Membranes were probed with respective antibodies.

### 2.6. Immunofluorescence microscopy and TUNEL assay

Cells were seeded at a density of 0.05 × 10^6^ cells per well in 24 well plates containing a round glass coverslip. After 24 hr, cells were serum starved for 12 hr. Then cells were cultured in 5% ND or 5% HFD serum-containing media for 48 hr. The protocol followed is described previously by Malvi P. et al., 2015 [13]. pH2AX antibody was used for immunofluorescence analysis by Zeiss confocal microscopy. Data were processed by ZEN desk software.

Terminal deoxynucleotidyl transferase dUTP nick-end labeling (TUNEL) assay in cells was performed as per the manufacturer of the TUNEL staining kit (556381) protocol. TUNEL staining kit (556381) was purchased from BD Biosciences (NJ, USA)).

### 2.7. Comet assay

Cells were seeded in 12 well plates at a density of 0.1×10^6^ cells per well and allowed to adhere for 24 hr, which was followed by serum starvation. Afterwards, ND or HFD serum-containing media was added and cells were cultured for 48 hr. Trypsinization of cells was followed by centrifugation at 2000 rpm for 2 min. The cell pellet was suspended in 100 µl of PBS. The cell suspension was mixed with melted 0.5% low melting agarose (LMA) and then spread on 1% normal melting agarose (NMA) coated dried slides. Cells were lysed in cold lysis solution for 2 hr and then incubated in electrophoresis buffer for 20 min in the dark. Electrophoresis was performed for 30 min at 300 mA in the dark. The neutralizing solution was added to slides for 15 min and DNA was stained with ethidium bromide (20 µg/ml, EtBr). The migration of DNA was observed using a fluorescent microscope (Olympus microscope, Tokyo, Japan), and images were captured. Data analysis and quantification were performed using the Open Comet plugin in Image J software.

### 2.8. Cell proliferation assay

Cells were seeded in 96 well plates (0.005×10^6^ cells per well) and allowed to adhere for 24 hr. These were serum starved for 12 hr followed by replacement with media containing serum derived from ND or HFD mice. At various time points 3-(4,5-dimethylthiazol-2-yl)-2,5-diphenyl tetrazolium bromide (MTT) was added and percent proliferation was determined as mentioned by Malvi P et al., 2015 [13].

### 2.9. Cell cycle analysis

Approximately 0.3×10^6^ cells/well were seeded in 6 well plates. Cells were incubated for 24 hr followed by 12 hr of serum starvation. After 48 hr of culture in 5% ND or 5% HFD serum-containing media, trypsinization was performed. Cells were washed with chilled PBS. Fixing of cells was performed with 70% chilled ethanol. Then cell pellet was washed twice with PBS, followed by treatment with RNase A (200 µg/ml) and incubated in the dark for 30 min. DNA staining dye propidium iodide (PI-50 μg/ml) was added and the cell suspension was transferred into FACS tube using cell strainer. Data was acquired by flow cytometer (FACS, Caliber) and analyzed using Cell Quest Pro Software.

### 2.10. RNA isolation and real-time PCR

Total RNA from cells was extracted using TRIZOL reagent as per the manufacturer’s (Invitrogen, CA, USA) protocol. RNA was then quantified by the spectrophotometric method using nanodrop. After extraction of total RNA, the first strand cDNA was synthesized from 2 µg RNA. Mouse ercc1, xrcc4, xrcc6, rad51, msh2, and 18s rRNA-specific primers were used for quantitative real-time polymerase chain reaction (qRT-PCR).

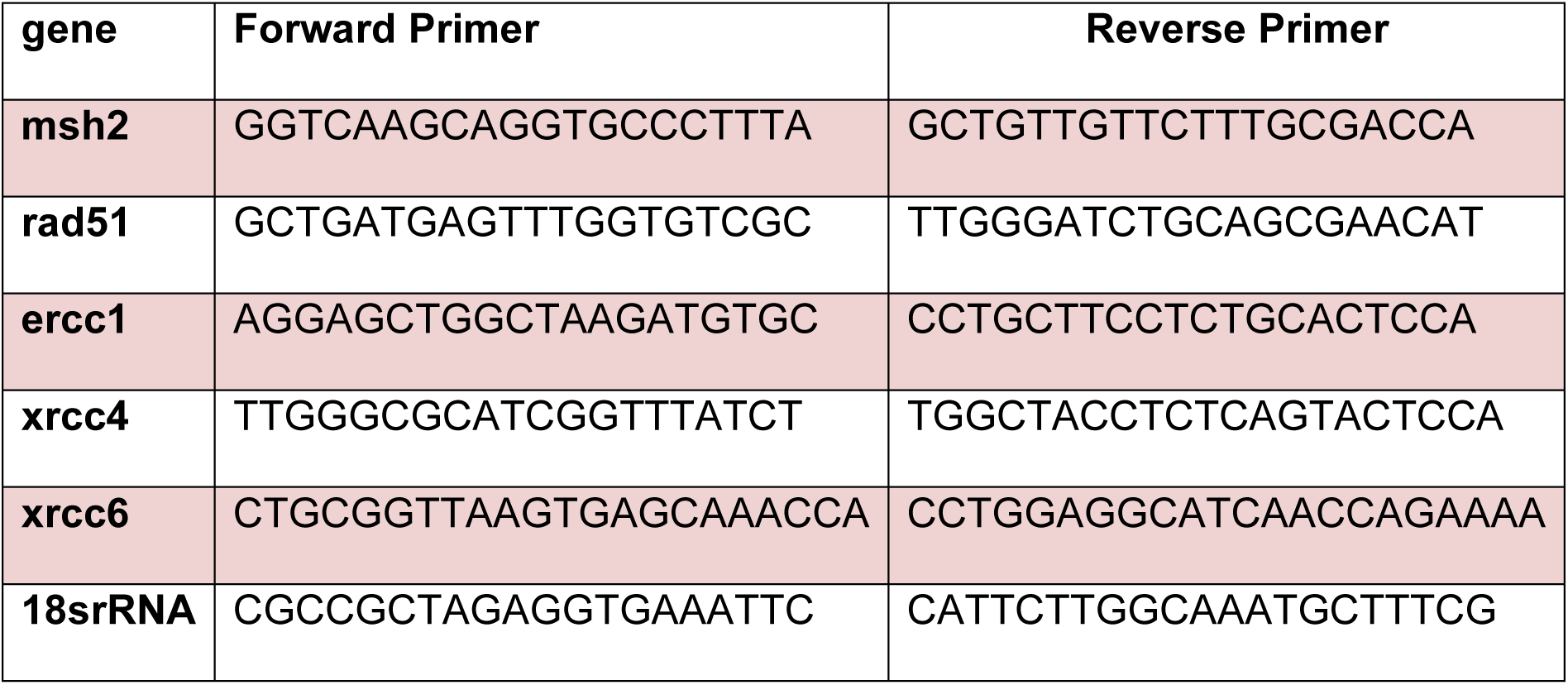

### 2.11. Statistical analysis

Graph Pad Prism 8 Software was used for statistical analysis (student’s two-tailed unpaired t-test) to compare values between experimental groups. Data was represented as mean± standard error of the mean (SEM). Bar represents values within wells of an experiment. The values of P<0.05 were considered statistically significant (*P<0.05, **P<0.01, ***P<0.001).

## 3. Results

### 3.1. HFD serum factors induce DNA damage in cells

An earlier report from our laboratory suggested that HFD-fed obese mice exposed to a carcinogen-azoxymethane exhibited an increased occurrence of colon polyps as compared to ND-fed control mice. As the metabolite of AOM causes mutation in crucial genes by alkylation of DNA, obese factors may contribute to aggravate DNA damage. To explore the role of obesity-associated serum factors on DNA damage *in-vitro*, mouse-derived colon cancer cells CT26 (p53 wild-type), MC38 (p53 mutated), and mouse-derived non-cancerous cells 3T3-L1 (p53 wild-type) were cultured in 5% ND or 5% HFD serum-containing media for 48 hr. It was observed that the fluorescence intensity of TUNEL staining in CT26 and 3T3-L1 cells cultured in HFD serum was increased as compared to ND serum (Figure 1A&B). No noticeable differences were detected in MC38 cells cultured in HFD and ND serum (Figure 1C). A specific assay commonly used to assess the extent of DNA damage is a comet assay. The percentage of DNA in the tail of comets, which is indicative of DNA damage was increased in CT26 and 3T3-L1 cells cultured in HFD serum compared to cells cultured in ND serum (Figure 1D). Also, at the molecular level, the expression of DNA damage marker pH2AX was increased in CT26 and 3T3-L1 cells cultured in HFD serum as checked by immuno-fluorescence and immuno-blot analysis (Figure 2A,B,D&E). Put together these results indicate that obesity-associated serum factors induce DNA damage in a cell type-dependent manner.

**Figure 1.**
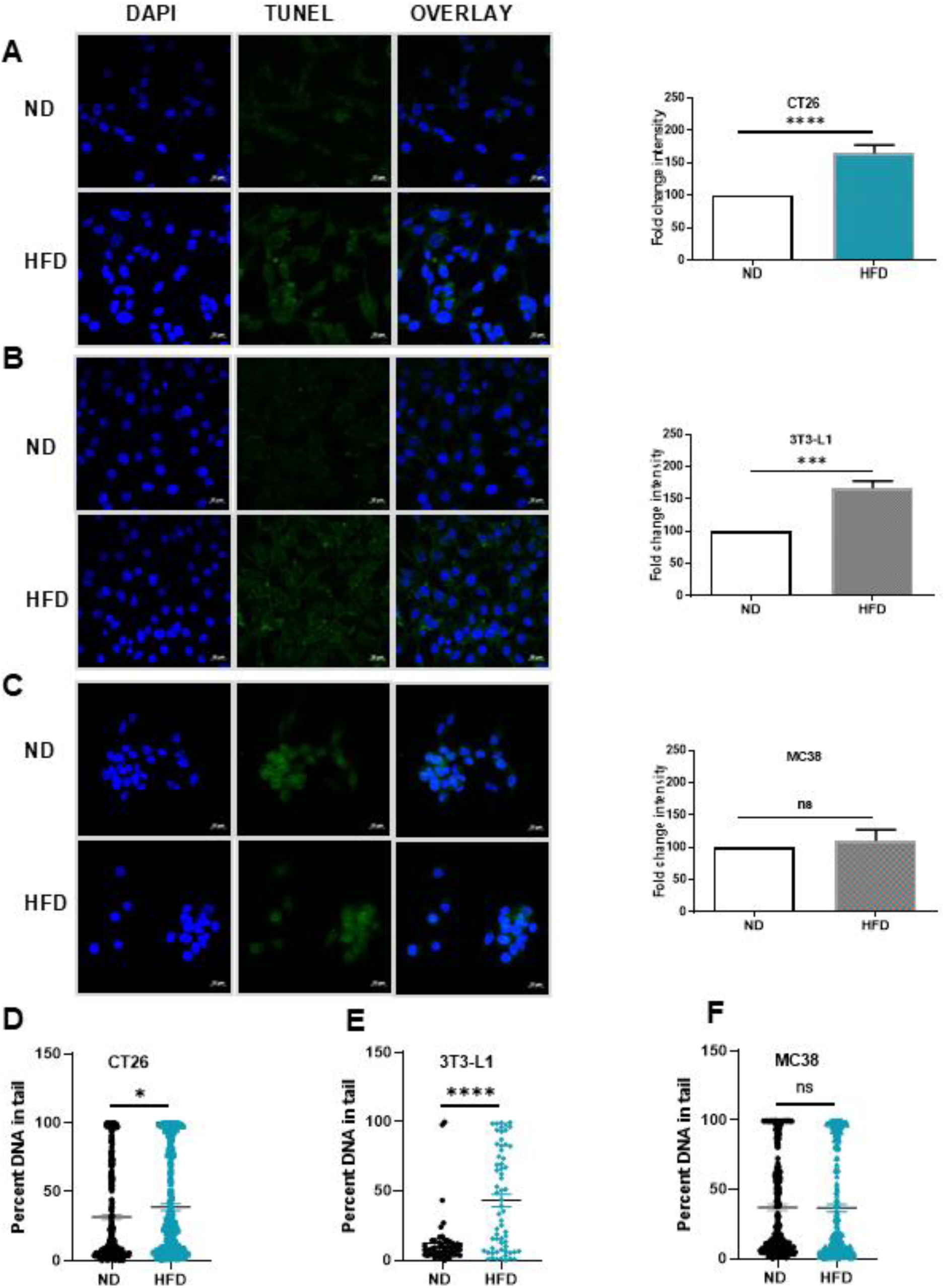
Analysis of the extent of DNA damage in cells cultured in ND and HFD serum. CT26, 3T3-L1, and MC38 cells were cultured in serum collected from ND and HFD-fed mice for 48 h.; TUNEL and comet assays were performed as per the protocol mentioned in the methods—fluorescent images were taken at 40X magnification by confocal microscope. The bar graph represents the quantification of fluorescence intensity in multiple fields of coverslip by using Image J software. (A) CT26 cells (B) 3T3-L1 cells, and (C) MC38 cells. Experiments were performed twice. (D, E&F) Phase contrast images of comets in CT26, 3T3-L1, and MC38 cells at 4X magnification. The bar graph represents the intensity quantification in multiple fields of the slide by using Image J software. The data represented value in avegare±SEM of three fields of a slide. The experiment was repeated twice.

**Figure 2.**
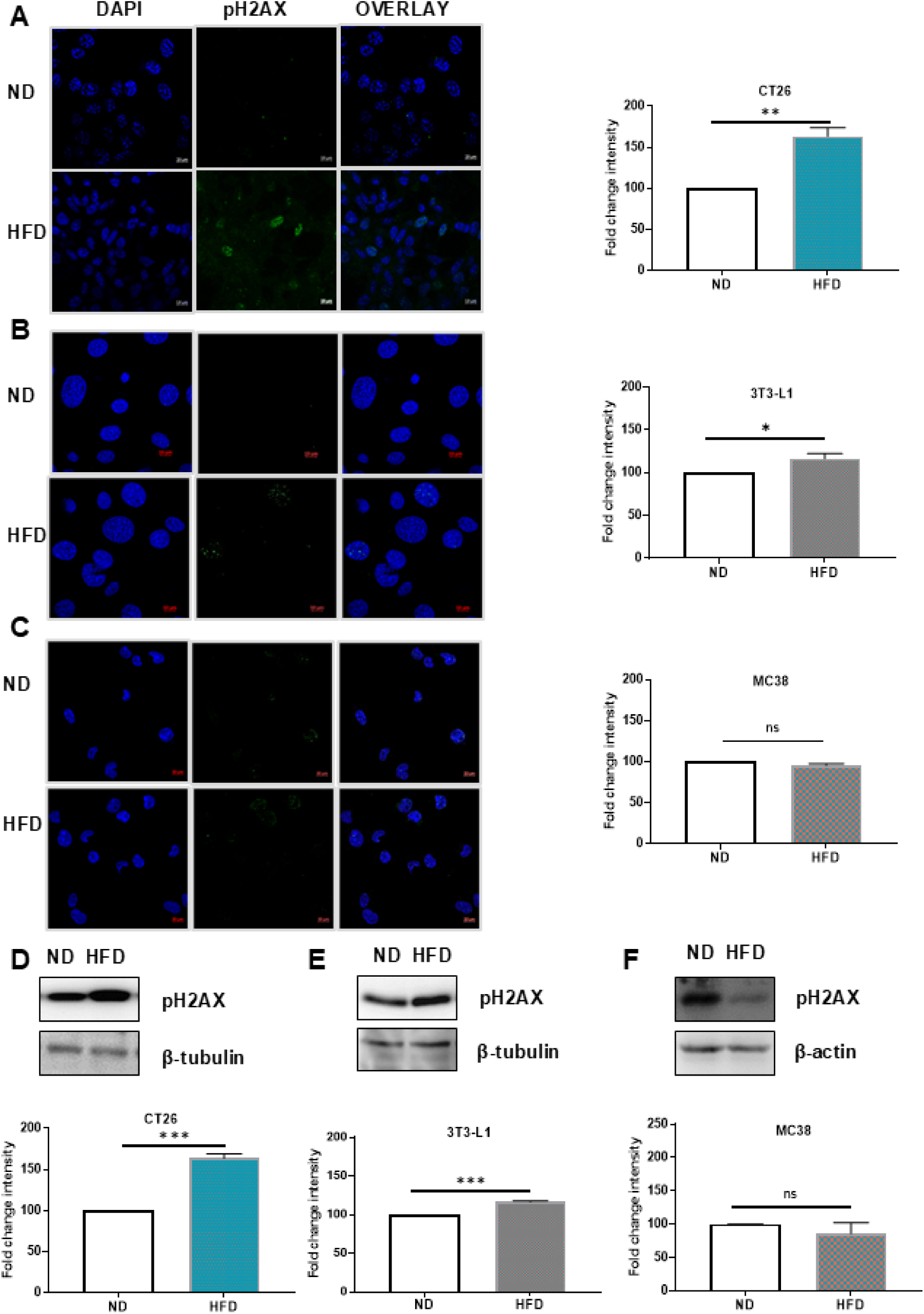
Induction of DNA damage marker in cells cultured in HFD serum. CT26, 3T3-L1, and MC38 cells were cultured in serum collected from ND and HFD-fed mice for 48 hr. Immunofluorescence and immunoblot analysis were performed as per the protocol mentioned in the methods. (A, B&C) Immunofluorescent images were taken at 40X (CT26 cells) and 60x (3T3-L1 and MC38) magnification by confocal microscope. The bar graph represents the quantification of fluorescence intensity in multiple fields of coverslip by using Image J software. Experiments were repeated twice. (D, E&F) Immunoblot analysis of pH2AX protein. Expression was evaluated from the whole cell lysate of CT26, 3T3-L1, and MC38 cells. β-tubulin/actin was used as a loading control. The bar graph represents the intensity quantification by using Image J software. The experiment was performed three times.

### 3.2. HFD serum alters DNA damage response

To further evaluate the effect of HFD serum on DDR in cells, the status of critical molecules of DDR such as P-p53ser15 and pchk2 were analyzed. Various stress-inducing factors stimulate multiple phosphorylation sites on p53. However, p53ser15 phosphorylation is one of the focal points for cellular stress response [18]. Another important DDR molecule chk2 is phosphorylated at Thr 68 by transducer kinases in response to DNA damage [19]. As indicated earlier, an increase in DNA damage was observed in cells cultured in obese serum and thus, the status of DDR molecules was probed. In CT26 and 3T3-L1 cells cultured in ND and HFD serum basal level expression of p53 was observed to be comparable. However, P-p53ser15 levels were significantly increased in HFD serum cultured cells compared to cells cultured in ND serum (Figure 3A,B,C&D). Additionally, the activated form of upstream mediator molecule chk2 was increased in CT26 cells cultured in HFD serum, indicative of activation of the DDR pathway (Figure 3E&F).

**Figure 3.**
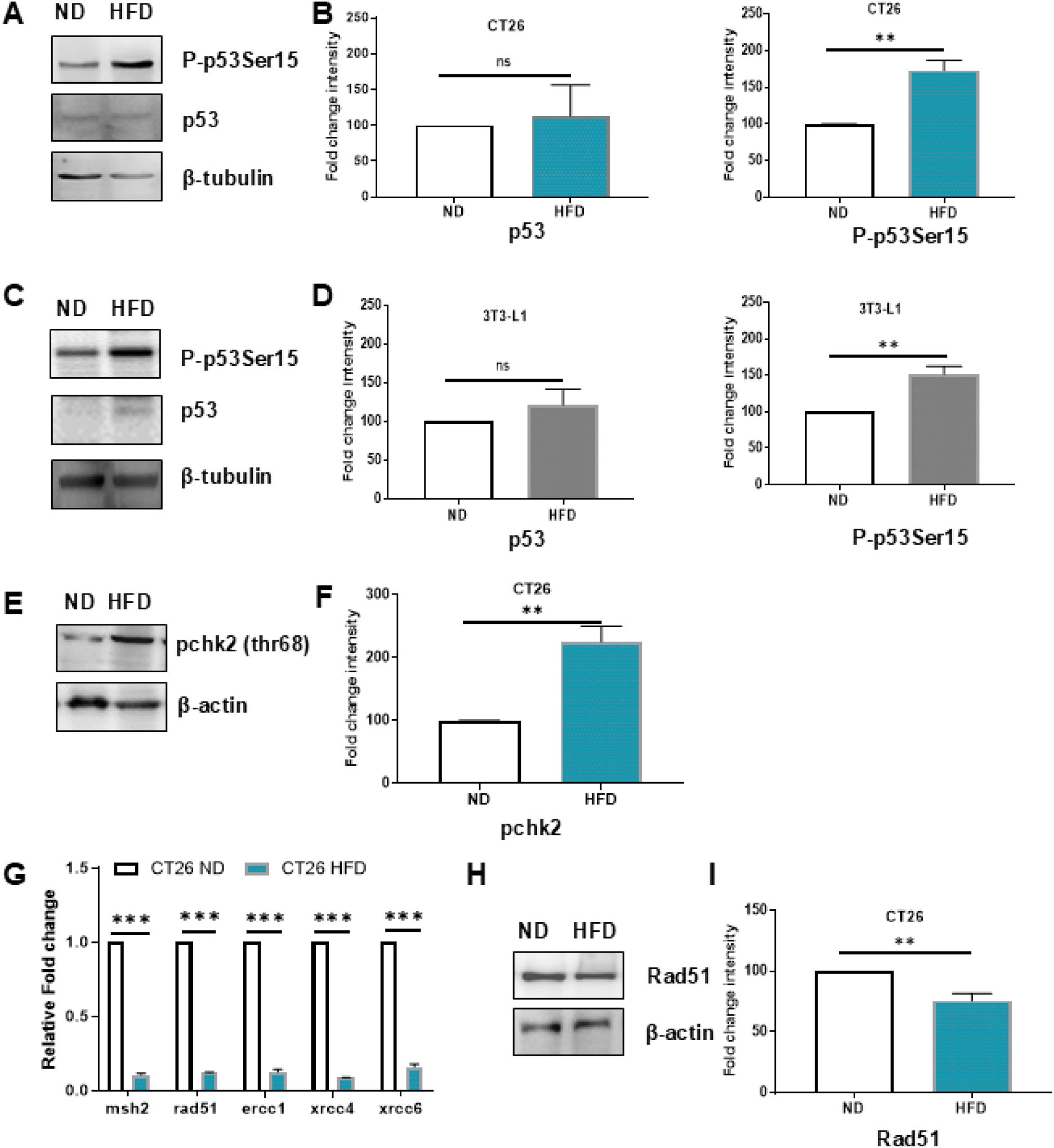
Molecular changes associated with DNA damage response in cells cultured in ND and HFD serum. CT26 and 3T3-L1 cells were cultured in ND and HFD serum for 48 hr; followed by an analysis of molecules involved in DNA damage response by immunoblotting and DNA repair mechanism by real-time PCR. Cells were cultured in ND and HFD serum, and immunoblotting for P-p53ser15 and p53 was performed in the whole cell lysate. β-tubulin was used as the loading control. Bar graph represents the intensity quantification by using Image J software. The experiment was performed three times (A&B) CT26 cells (C&D) 3T3-L1 cells.The experiment was performed threetimes. (E) CT26 cells were cultured in ND and HFD serum, and immunoblotting for pchk2 was performed in the whole cell lysate β-actin was used as the loading control. (F) The bar graph represents the intensity quantification by using Image J software. The data represents value from three independent experiments. (G) CT26 cells were cultured in ND and HFD serum and real-time PCR was performed using specific primers for msh2, rad51, ercc1, xrcc4, and xrcc6 mRNA. Experiment were done in triplicate and performed twice.(H) CT26 cells were cultured in ND and HFD serum, and immunoblotting for Rad51 was performed in the whole cell lysate. β-actin was used as the loading control. (I) The bar graph shows the intensity quantification by using Image J software. A similar experiment was performed three times.

An efficient repair system restores damage in DNA and maintains the integrity of the genome. Further the status of DNA repair molecules was evaluated in HFD cultured cells. Interestingly, the mRNA levels of msh2, rad51, ercc1, xrcc4, and xrcc6 were significantly decreased in CT26 cells cultured in HFD serum (Figure 3G). Also, the protein level of an important repair molecules Rad51 was analyzed by immunoblotting. In CT26 cells cultured in HFD serum Rad51 level were significantly decreased (Figure 3H). Similar trends were also observed in 3T3-L1 cells (Figure S1). These results indicate that HFD serum alters DDR and downregulates repair molecules.

### 3.3. AOM/DSS causes increase in polyp occurrence in the colon of *ATM-/-* mice

There are three apical kinases ATM, ATR, and DNA-PKCs which phosphorylate H2AX at ser139. To address the effect of functional implication of absence/presence of DDR molecule in initiation of carcer *ATM+/+* and *ATM-/-* mice were used in present study. *ATM-/-* mice exhibit altered metabolic parameters such as glucose intolerance, and lower serum adiponectin levels together with reduced accumulation of subcutaneous adipose tissue suggestive of a systemic effect of ATM molecule [20]. The mutations in *ATM* gene also promote genomic instability and predispose cells to transformation [21]. We first investigated the functional implication of ATM in the context of cancer occurrence. Thus, ND fed *ATM+/+* and *ATM-/-* mice were administered with AOM/DSS to induce polyp formation in colon as per the experimental layout (Figure 4A). Interestingly, the incidence of polyp formation was higher in *ATM-/-* (3/5) mice as compared to *ATM+/+* (1/3) mice (Figure 4F). This result indicates that the non-functionality of ATM increases the risk of carcinogen-induced initiation of colon cancer in mice. Also, it was observed that in AOM/DSS administered *ATM-/-* mice the levels of serum triglyceride (TG) and cholesterol were increased as compared to *ATM+/+* whereas no significant differences in the weight of the animals were observed. (Figure 4C,D&E). Also in another set of experiments in HFD-fed, *ATM+/+* and *ATM-/-* mice no observable differences were detected in the weight of the mice, serum TG and cholesterol levels (Figure S2).

**Figure 4.**
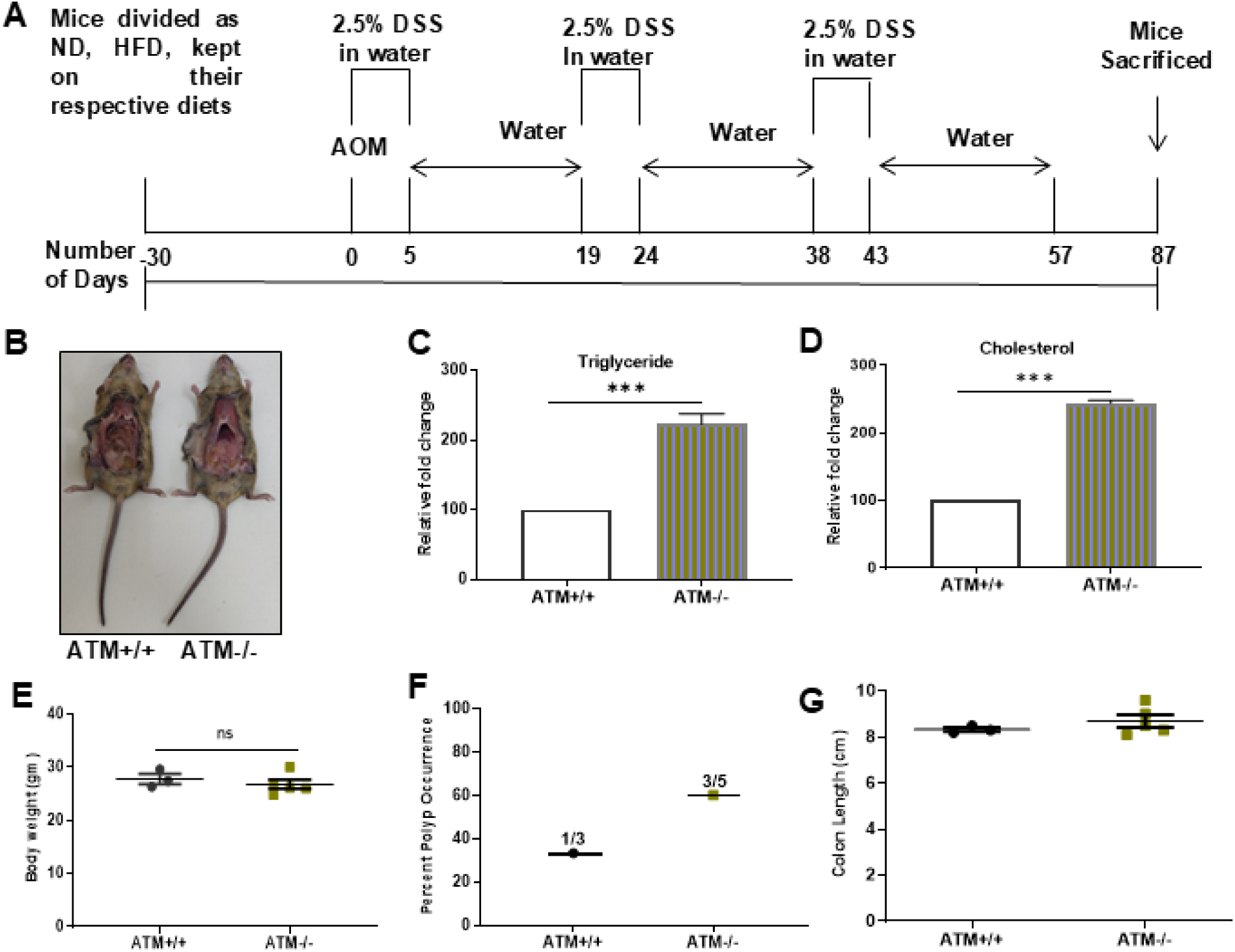
Effect of absence of ATM, a DDR-associated molecule. (A) *ATM+/+* and *ATM-/-* mice were administered with AOM/DSS for polyp formation in the colon. Polyps were developed by injecting AOM (10 mg/kg body weight) and followed by DSS (2.5%) in drinking water. Experimental design for AOM/DSS induced colorectal carcinogenesis. (B) Photographs of *ATM+/+* and *ATM-/-* mice were taken at the end of the experiment. (C) Body weight of *ATM+/+* and *ATM-/-* mice. (D) Triglyceride levels in serum of *ATM+/+* and *ATM-/-*. (E) Cholesterol levels in serum of *ATM+/+* and *ATM-/-*. (F) Percentage of polyp occurrence in *ATM+/+* and *ATM-/-* mice. (G) Colon length of *ATM+/+* and *ATM-/-* mice.

### 3.4. HFD serum-associated factors increase survival and pH2AX level in colon cancer cells

Previous reports from our laboratory have shown that HFD serum and obesity-associated adipokines promote proliferation of cancer cells [17]. Hence, we hypothesized that factors present in HFD serum, though inducing DNA damage, contribute towards the survival of cancer cells. Therefore, to investigate cell survival, HCT116 +/+ (p53 wild-type) were used. One of the survival molecules is survivin, which is the smallest member of the inhibitor of apoptosis protein (IAP). Survivin also regulates cell cycle by binding to microtubules and chromosomes, thereby favoring cell survival [22]. The survivin protein levels were found to be increased in HCT116+/+ cells cultured in HFD serum (Figure 5A&B). To further asscetain the survival phenotype of cells, the levels of phosphorylated ser15, ser46, and ser20 of p53 were analyzed. Earlier reports indicate that an unaltered level of ser46 and an increased level of ser15 and ser20 of p53 skew the fate of cells toward survival [23]. Also, the levels of phosphorylated ser15 and ser20 were elevated whereas, the levels of phosphorylated ser46 remained unaltered in HCT116 cells cultured in HFD serum compared to ND serum (Figure 5C). Moreover, decrease in apoptotic population was detected in the cells cultured in HFD serum (Figure S3). Collectively, these results are indicative of potential molecular events contributing to cell survival despite increased DNA damage.

**Figure 5.**
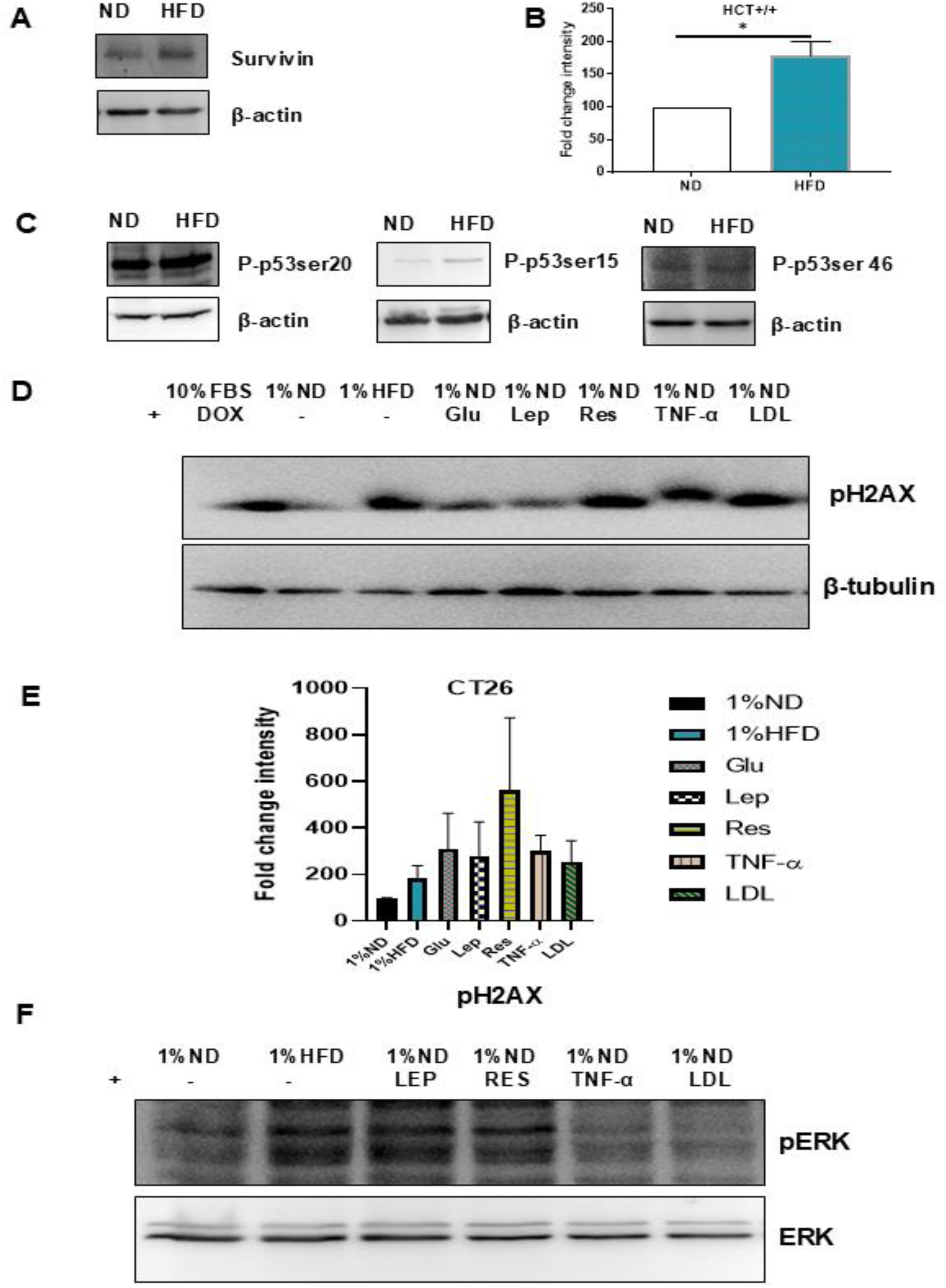
HFD serum-associated factors affect molecules associated with survival and DNA damage in colon cancer cells. HCT116+/+ were cultured in ND and HFD serum for 48 hr; and cell lysates were used for immunoblot analysis. (A) Survivin levels in whole cell lysate. (B) Bar graph represents the intensity quantification of survivin in HCT+/+ cells by using Image J software. (C) Analysis of P-p53ser20, P-p53ser15 and P-p53ser46 in whole cell lysate. (D) Analysis of pH2AX in CT26 cells cultured in 1% ND, 1% HFD, 1%ND and glucose, leptin, resistin, TNF--α, and LDL. (E) Fold change in the intensity of pH2AX in CT26 cells cultured in 1% ND, 1%HFD, 1%ND and glucose, leptin, resistin, TNF-α, and LDL. (F) Analysis of pERK in CT26 cells cultured in 1% ND, 1% HFD, 1%ND, and leptin, resistin, TNFα and LDL.

In obese condition, the status of serum biochemical parameters such as adipokines, glucose, and cholesterol (LDL/HDL) get altered. Therefore, to elucidate which of the serum-associated factors might contribute to increase in pH2AX level, CT26 cells were cultured in ND or HFD, and ND containing14 mM glucose or 100 ng/ml mLeptin (mouse) or 100 ng/ml mResistin or 100 ng/ml mTNF-α or 50 µg/ml hLDL (human) or 50 µg/ml hHDL. Interstingly, In the cells cultured with media containing mLeptin, mResistin, mTNF-α, and hLDL, an increase in pH2AX level was detected (Figure 5D&E). Earlier data from our laboratory reported that leptin treatment causes ERK activation and also induces cell proliferation [17]. On similar lines, an increase in ERK level was observed in leptin and resistin-treated cells (Figure 5F).

### 3.5. HFD serum promotes proliferation of cells

Increase in DNA damage and altered DDR together with the survival of colon cancer cells cultured in HFD serum led us to explore the parameters of growth of cells. Interestingly, CT26 and 3T3-L1 cells cultured in HFD serum for 48 hr and 72 hr proliferated rapidly as compared to cells cultured in ND serum. No change in growth of MC38 cells was detected at 24 hr, 48 hr, and 72 hr (Figure 6A,B&C). Additionally, HFD serum differentially influences the proliferation rate of HCT116+/+, HCT116-/-, Huh7, and CHO cells (Figure S4). Moreover, cell cycle analysis by FACS indicated an increase in the percentage of CT26 and 3T3-L1 cells in the S phase and decrease in the ratio of G0G1 to S population, upon culturing in HFD serum (Figure 6D,E&F; Figure 6G,H&I). Also, colony formation and soft agar assay indicate proliferative phenotype of CT26 and 3T3-L1 cells (Figure S5 and S6). However, no differences were noticed in cell cycle phases of MC38 cells cultured in ND and HFD serum.

**Figure 6.**
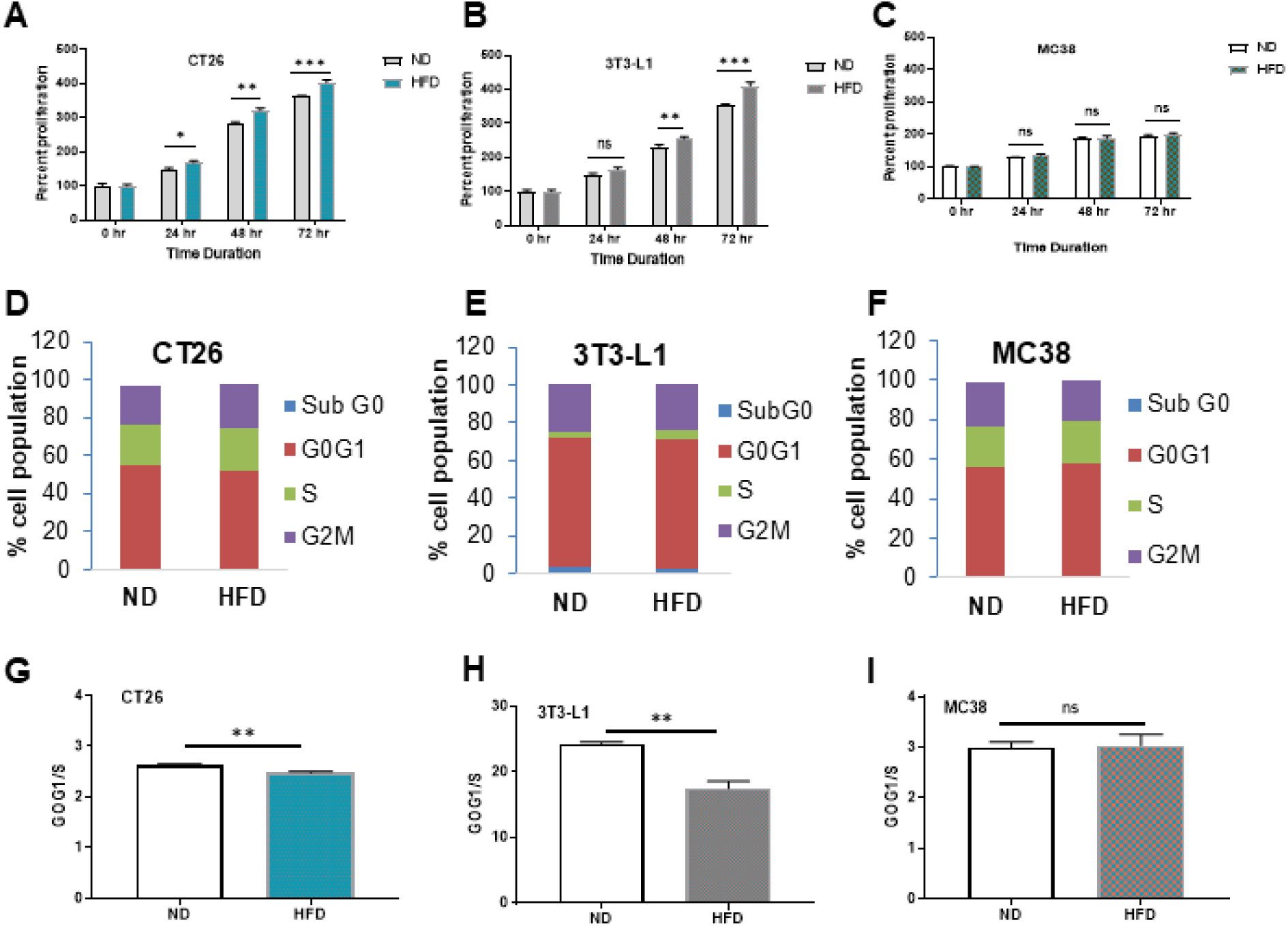
Analysis of cell proliferation. CT26, 3T3-L1, and MC38 cells were cultured in ND and HFD serum; MTT and FACS analysis was performed as per protocol mentioned in methods. (A, B&C) CT26, 3T3-L1, and MC38 cells were cultured in ND and HFD serum-containing media for different time points i.e. 24 hr, 48 hr, and 72 hr and percent cell proliferation was calculated through MTT assay. (D, E&F) Bar graphs represents % cell population in different phases of the cell cycle of CT26, 3T3-L1, and MC38 cells; the population of subG0, G0G1, S, and G2M cell population. (G, H&I) Bar graphs represents the ratio of G0G1 to S (G0G1/S) of CT26, 3T3-L1, and MC38 cells cultured in ND and HFD serum.

### 3.6. Polyp occurrence was increased in AOM/DSS-administered HFD mice

The metabolite of azoxymethane causes alkylation of bases in DNA leading to mutations in critical signaling molecules such as β catenin, TGF β, and K-ras [24]. Moreover, obesity has been linked to the occurrence, and proliferation of cancer cells [12] [17]. To evaluate the consequence of AOM treatment on HFD-fed mice, twenty four C57BL/6J mice were divided into ND and HFD groups. Mice were fed with respective diets for a month. The AOM/DSS treatment cycle was followed as per the details mentioned in Figure 4H. After the completion of treatment cycle, four mice from each group were monitored for survival analysis (Figure 7G). Also, to analyze the polyp formation, eight mice from each group were sacrificed. Interestingly, in HFD-fed mice, polyp occurrence and the number of polyps per colon were more suggesting that obesity aggrevates the risk of cancerous polyp formation upon AOM administration compared to ND-fed mice (Figure 7C,D&E). The length of colon of HFD-fed mice was significantly reduced than ND fed mice (Figure 7B). The methylene blue staining of colon polyps is indicative of dysplastic phenotype. Moreover, detection of increased expression of β-catenin, PCNA, and pH2AX in polyp lysates compared to normal tissue is indicative of neoplastic phenotype (Figure S7). However, survival data of the AOM-administered ND and HFD mice group revealed prolonged survival of HFD mice.

**Figure 7.**
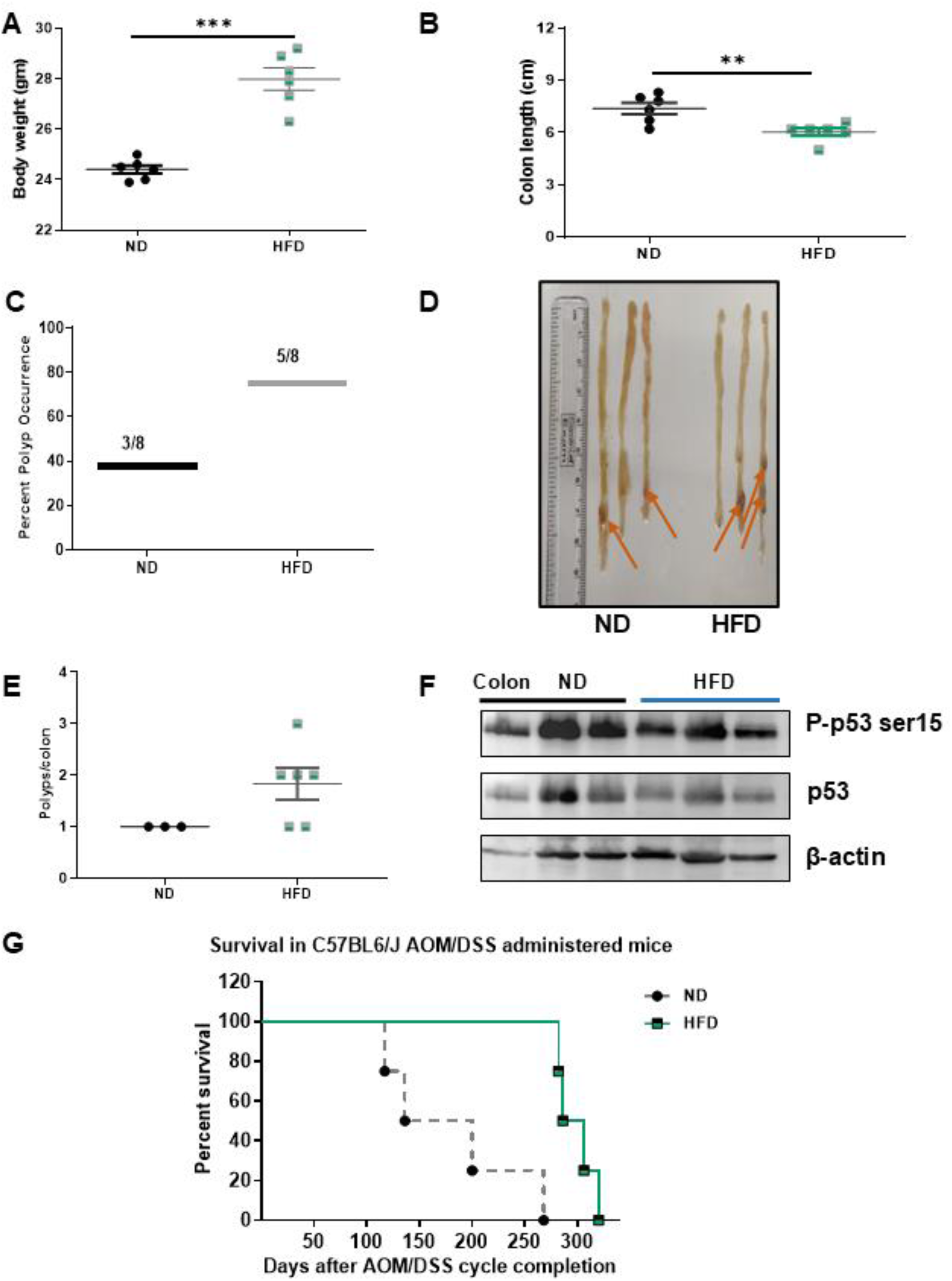
AOM/DSS induced colon cancer in ND and HFD-fed C57BL/6J mice. Chemically induced colon polyps were developed by injecting AOM (10mg/kg body weight) and followed by DSS (2.5%) in drinking water. (A) Mice body weight recorded before AOM administration. (B) Colon length of ND and HFD fed mice. (C) Percentage of polyp occurrence in ND and HFD-fed mice. (D) Photograph of polyp bearing colons. (E) No. of polyps per colon in each group. (F) Immunoblot analysis of P-p53ser15 and p53 in colon tissue lysate. β-actin was used as loading control. (G) Survival curve of AOM-administered ND and HFD mice. After completion of the AOM/DSS cycle, 4 mice from each group were randomly selected for survival analysis which may or may not have polyps in the colon. The graph shows the percent survival of AOM/DSS administered ND and HFD mice.

## 4. Discussion

Several clinical as well as pre-clinical studies have reported increase in DNA damage in various cell types under obese conditions [6], [25]–[27]. The present study demonstrates that HFD serum-associated factors contribute to increased DNA damage and altered DNA damage response, in cultured cells. To our knowledge, this is the first study on the obesity-altered DNA damage response specifically in colon cancer cells.

Our data indicate that DNA damage occurrence is cell type-dependent, with certain cells exhibiting increased DNA damage in obese conditions, while others don’t demonstrate (Figures 1 and 2). This suggests a potential role of the DNA damage response (DDR) machinery in regulating these differences. To investigate this, we analyzed the expression of DNA damage and repair-related molecules. Our results show an increase in DNA damage markers (P-p53Ser15 and pChk2) and a decrease in DNA repair proteins (MSH2, RAD51, ERCC1, XRCC4, and XRCC6) in CT-26 cells cultured with HFD serum compared to ND serum (Figure 3).The simultaneous activation of DDR and downregulation of DNA repair proteins may appear contradictory; however, cancer cells are inherently genetically instable. Under obese conditions, the reduced levels of repair proteins are insufficient to effectively repair DNA damage, leading to sustained DDR activation. Cancer cells have the advantage that despite the presence of DNA damage, cancer cells continue to proliferate rapidly.

Clinical reports also suggest a positive link between DNA damage and adiposity [6], [18], [29]. However, most of these reports have assessed DNA damage in normal cells rather than cancer cells. Notably, an article from David A. Cavazos et al., reported an increase in DNA damage in prostate cancer cells cultured in HFD serum. Results revealed that circulatory factors of serum from HFD mice increase the level of pH2AX, elevate ROS generation, and facilitate proliferation of prostate cancer cells [30].

To assess linked molecular events in present study, we evaluated the role of ATM. Using an *in-vivo* AOM/DSS-induced colon cancer model in ATM+/+ and ATM−/− mice, we observed increased polyp formation in ATM-deficient mice. These findings suggest that the absence of DDR molecule-ATM has role in increased risk of cancer occurance (Figure 4). Earlier reports have reported that ATM deficiency accelerates carcinogenesis in KRAS-induced pancreatic cancer model [28].

In obesity, serum parameters such as adipokines, cholesterol and triglycerides levels are altered [13]. To mimic whether altered serum levels of these factors have influence on molecular events, cells were cultured in the presence of adipokines and cholesterol. Our data indicate that HFD serum and its components, leptin, resistin, TNFα and LDL induce DNA damage (Figures 5D&5E). Also, leptin and resistin activates ERK, which may be involve in increased proliferation of cells (Figures 5F). Moreover, the role of individual factors associated with obesity such as cholesterol, glucose, and specific adipokines like leptin, resistin, TNF-α, adiponectin, and visfatin, in the proliferation of various cancer cell has been reported [17], [36]. Importantly, on clincal front, a review of available data has lead to validating the crucial role of obesity-associated factors in tumor initiation and progression [37]. A published article from our laboratory suggested the specific role of leptin in proliferation of cancer cells through involvement of ERK [17]. These findings highlight the differential effects of HFD serum and its components on cell proliferation and DNA damage. In present study the increased polyp formation in HFD-fed mice and elevated DNA damage in cultured cells may be attributed to the downregulation of DNA repair, potentially mediated by specific serum components, which warrants further investigation.

Previously, our laboratory reported that HFD-fed mice exhibit higher carcinogen-induced polyp formation compared to leptin receptor knockout (ob/ob) mice. The reduced polyp frequency in ob/ob mice suggests a direct role of a combination of adipokines in tumorigenesis. However, whether this effect is mediated through DNA damage or DNA repair mechanisms remains to be elucidated.

Recent studies highlighted the role of leptin induced p53 in breast cancer cells under obese phenotype [29]. However, more than 60% of colon cancers possess mutations in the p53 gene [30]. Therefore, colon cancer cells with varying p53 status were selected in this study. At the genetic level the mouse-derived colon cancer cells CT26 and MC38 have differences in the p53 status and in the functionality of leptin receptor. CT26 cells have functional and wild-type p53 whereas MC38 cells have mutant p53 and leptin receptors is mutated [17], [31]. For comparative investigations, the mouse-derived pre-adipocyte 3T3-L1 cells were utilized as these are non-cancerous, possess functional leptin receptors [32] and wild-type p53 [33]. By culturing the above-mentioned cells in ND and HFD serum, it was obseeved that the proliferation rate of CT26 and 3T3-L1 cells cultured in HFD serum was increased as compared to cells cultured in ND serum. However the the proliferation rate of MC38 cells was not influenced by presence of HFD or ND serum and therefore remained similar (Figure 6). It is likely that studies exploring the relationship of the proliferation of cancer cells under obese environment with p53 molecule at center would be helpful in delineating the mechanistic link between obesity and cancer.

As for relationship between obese phenotype and carcinogen effect *in-vivo*, AOM/DSS was administered to ND and HFD mice. It was observed that 75% of obese mice developed polyps in the colon, which is almost double than the perecentage of animals developing polyps incontrol group (37.5%) (Figure 7C). This finding is in agreement with the earlier reports [17]. Interestingly, it was observed that in mice with administered with AOM/DSS, HFD mice survived for a longer durtaion than ND mice (Figure 7G and Figure S7). This obeservation is in line with propostion made by Banack et al. who have comprehensively reviewed the articles in the context of obesity paradox. Also, it has been indicated that weight loss due to the illness is one of the paramount factor for normal-weight individuals, as compared to obese individuals, which likely is a contributory factor for survival benefit in different disease conditions [34].

Taken together this study paves the way for. investigations on the role of obesity-altered DDR as a contributory factor towards rapid proliferation of colon cancer cells. It points towards the important role of obesity in influencing nutritional and biochemical factors in cancers. However, the role of obesity-induced chronic inflammation, increased oxidative stress, altered metabolism, and immune response towards cancer development cannot be underestimated.

### Conclusion

The present study demonstrates that HFD serum factors induce DNA damage in cancerous and noncancerous cells *in-vitro*. It also causes alterations in DDR by increasing the levels of pH2AX, phosphorylated p53, chk2, and concurrently down-regulating the repair molecules. Remarkably, the induction of pH2AX is followed by a DDR induction that would either lead to cellular senescence or apoptosis. Instead, HFD serum promotes cellular viability. Together, the orchestrated deregulation of DDR and survival pathway supports the growth of damaged cells, likely predisposing them to additional transformations and proliferation.

## Acknowledgments

The authors thank Dr. S.C. Mande, and late Dr. Mohan R Wani, former Directors, BRIC-NCCS, Pune, India for being very supportive and giving all the encouragement to carry out this work. B.D. and F.B. thank University Grants Commission (UGC), New Delhi, India; H.Y. thanks Council for Scientific and Industrial Research (CSIR), India. The support from Experimental Animal Facility (EAF), Confocal Microscope Facility, Fluorescence Activated Cell Sorter (FACS) Facility, Central Instrumental Facilities, Technical Officer Dr Varsha Shepa**l,** and other group members are also duly acknowledged.

## Authors’ contributions

M.K.B., and B.D. conceptualized and designed the experiments, and wrote the manuscript; B.D. and H.Y. performed the experiments and analyzed the data with contributions coming from F.K.B. All the authors read, reviewed, and edited the manuscript.

## Ethics approval and consent to participate

All animal experiments have been performed as per the guidelines of the committee for the purpose of control and supervision of experiments on animals (CPCSEA), Government of India, and after obtaining the permission of the Institute’s Animal Ethics Committee (IAEC) (IAEC/2018/B-335 and IAEC/2018/B-336).

## Note

This work was carried out for the partial fulfillment of a Ph.D. thesis (of B.D.) submitted to Savitribai Phule Pune University, Pune, India.

## Additional Information

The dataset supporting the conclusions of this article is included within the article and its supplementary file (Figures S1 to S7).

## Abbreviations Used

RIPA: Radioimmunoprecipitation assay buffer
DDR: DNA damage response
ND: normal diet
HFD: high fat diet
AOM: azoxymethane
DSS: dextran sodium sulphate salt (35-50 KDa)
ATM: ataxia-telangiectasia
BMI: body mass index
IGF: insulin-like growth factor
8-OHdG: 8-hydroxy 2′-deoxy-guanosine
DMEM: dulbecco’s modified eagles medium
RPMI: roswell park memorial institute
EAF: experimental animal facility
CPCSEA: purpose of control and supervision of experiments on animals
IAEC: institutional animal ethics committee
PBS: phosphate buffer saline
TUNEL: terminal deoxynucleotidyl transferase dUTP nick-end labeling
LMA: low melting agarose
NMA: normal melting agarose
EtBr: ethidium bromide
MTT: 3-(4,5-dimethylthiazol-2-yl)-2,5-diphenyl tetrazolium bromide
PI: propidium iodide
qRT-PCR: quantitative real-time polymerase chain reaction
SEM: standard error of the mean
*BRCA*: breast cancer gene
IAP: inhibitor of apoptosis protein
LDL: low-density lipoprotein
HDL: high-density lipoprotein
TNF-α: tumor necrosis factor α

