## Supplemental Data 1 for "Obese serum factors aggravate DNA damage, alter DNA damage response, and promote proliferation in colon cancer cells"

**Supplementary figures:**

**Supplementary figure S1**

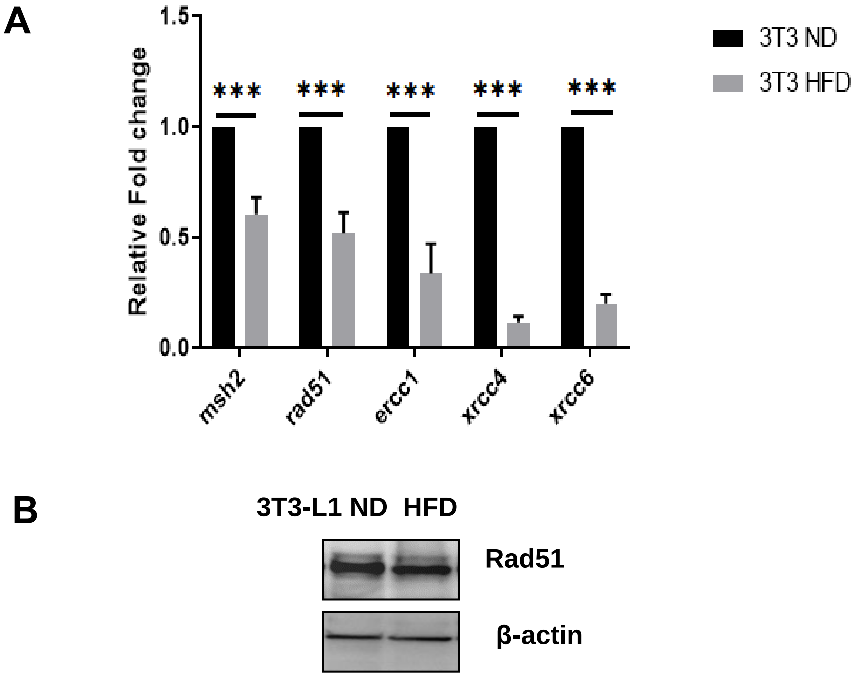


**Figure S1: Reduced expression of DNA repair molecules in 3T3-L1 cells cultured in HFD serum.**

(A) 3T3-L1 cells were cultured in ND and HFD serum and real-time PCR was performed using specific primers for msh2, rad51, ercc1, xrcc4, and xrcc6 mRNA. The experiment was done in triplicate. Relative transcript levels of repair molecules in 3T3-L1 cells. (B) 3T3-L1 cells were cultured in ND and HFD serum, and immunoblotting for Rad51 was performed in the whole cell lysate. β-actin was used as the loading control. The experiment was done in duplicate.

**Supplementary figure S2**

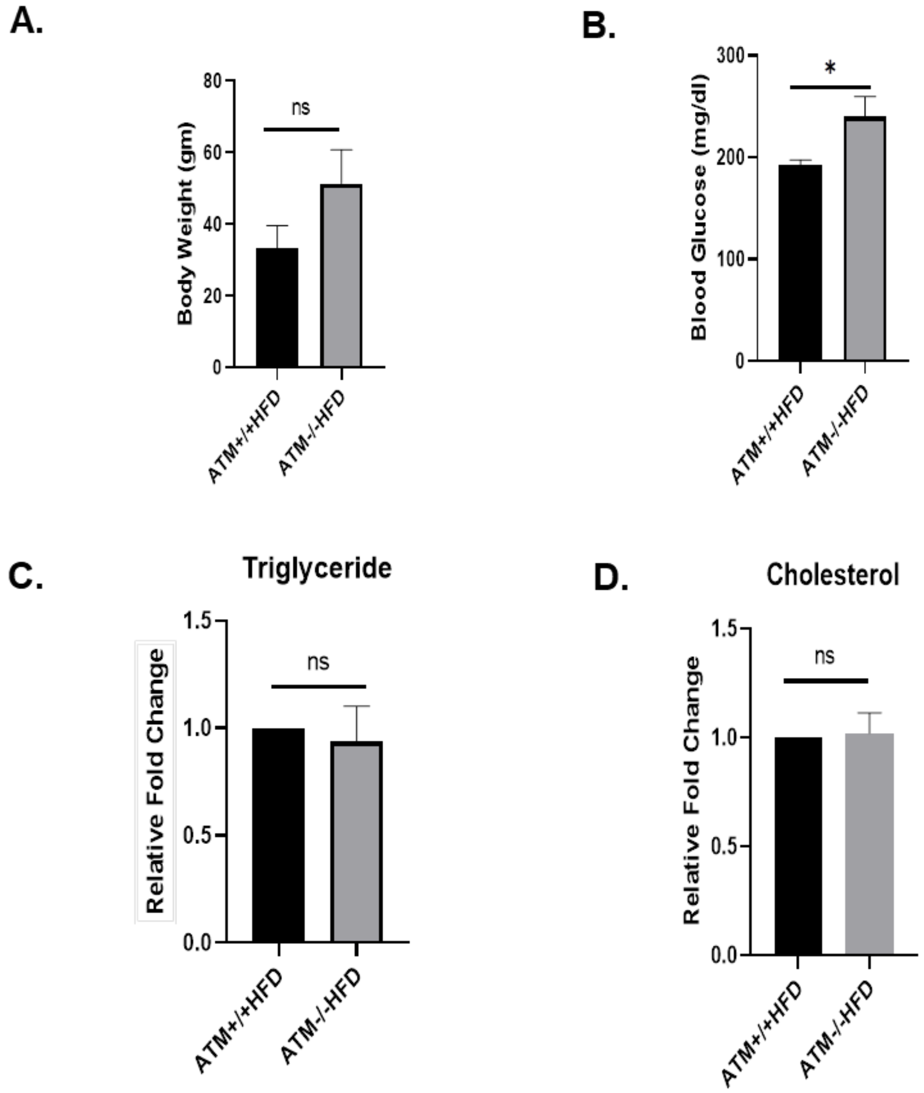


**Figure S2: Effect of HFD on serum parameters and body weight of *ATM+/+* and *ATM-/-* mice.**

129S6/SvEvTac-ATM(tm1Awb)/J *ATM*+/+ and 129S6/SvEvTac-ATM(tm1Awb)/J *ATM-/-* obese mice were developed by feeding HFD. *ATM* +/+ (n=3) and *ATM-/-* (n= 3) were grouped into *ATM* +/+ HFD and *ATM-/-* HFD groups and kept on respective diets for 5 months. *ATM+/+* and *ATM-/-* mice were fed HFD for 5 months. After that, blood was collected and serum parameters were analyzed (A) The bar graph represents body weight of *ATM+/+* HFD and *ATM-/-* HFD mice (B) The bar graph represents blood glucose levels in *ATM+/+* HFD, and *ATM-/-* HFD mice (C) The bar graph represents fold change in serum triglyceride levels of *ATM-/-* HFD mice (D) The bar graph represents fold change in serum cholesterol levels of *ATM-/-* HFD mice.

**Supplementary figure S3**

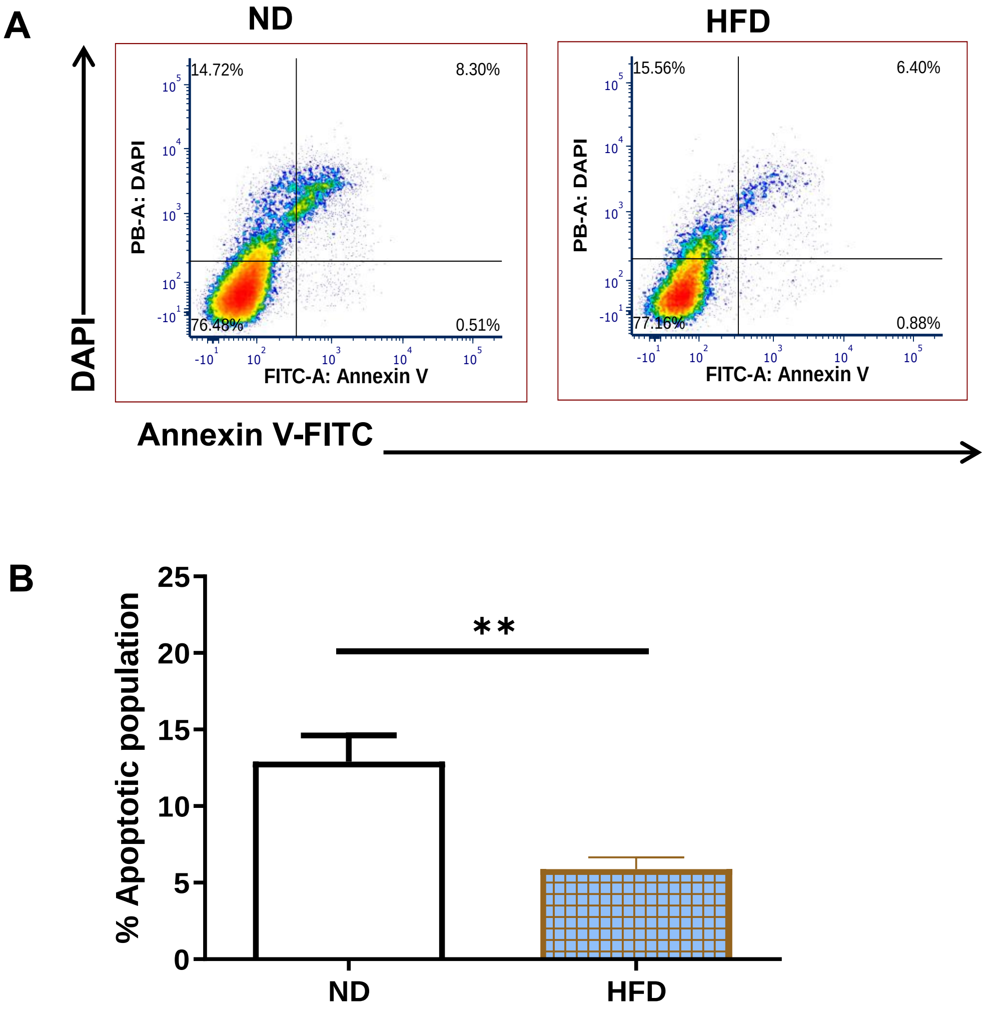

C

D

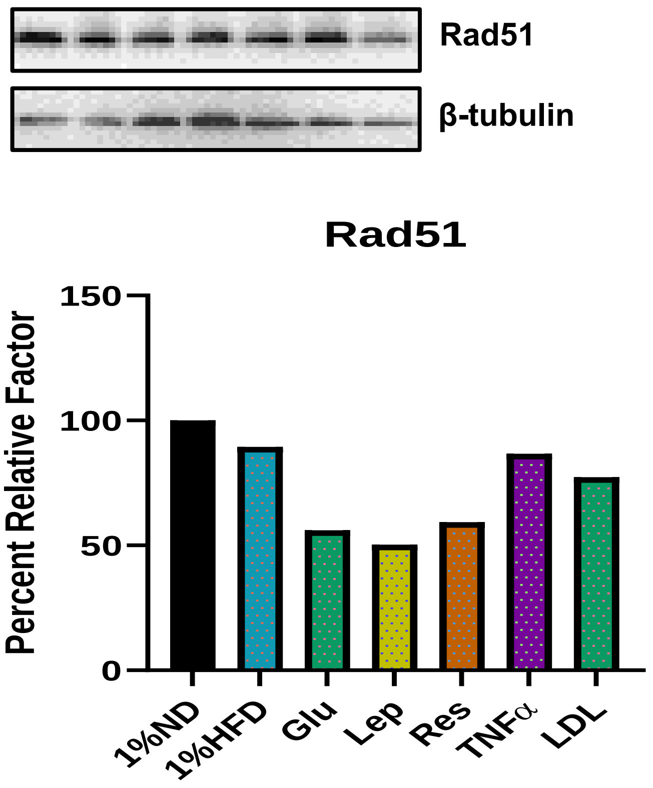


**Figure S3: Effect of obese serum on apoptosis of colon cancer cells.**

CT26 cells were seeded in 6 well plates at a density of 0.3x$10^6$ cells/well and allowed to adhere for 24 hr. After that cells were starved for 12 hr followed by culturing cells in ND or HFD serum-containing media for 48 hr. At the end of the experiment, the cells were trypsinized, washed with ice-cold PBS and stained with Annexin V/FITC in binding buffer (10 mM HEPES/NaOH, 40 mM NaCl, 2.5 mM $CaCl_2$) for 15-20 min in the dark at 37 ºC. (A) Dot plot showing different populations of live, early apoptosis, late apoptosis, and necrotic CT26 cells cultured in ND and HFD serum. (B) Bar graph showing total apoptotic population (early and late apoptosis) in ND and HFD. The experiment was performed in triplicates. (C) Analysis of Rad51 in CT26 cells cultured in 1% ND, 1% HFD, 1%ND and glucose, leptin, resistin, TNF-α, and LDL. (D) Fold change in the intensity of Rad51 in

CT26 cells cultured in 1% ND, 1%HFD, 1%ND and glucose, leptin, resistin, TNF-α, and LDL. This experiment was performed only once.

**Supplementary figure S4**

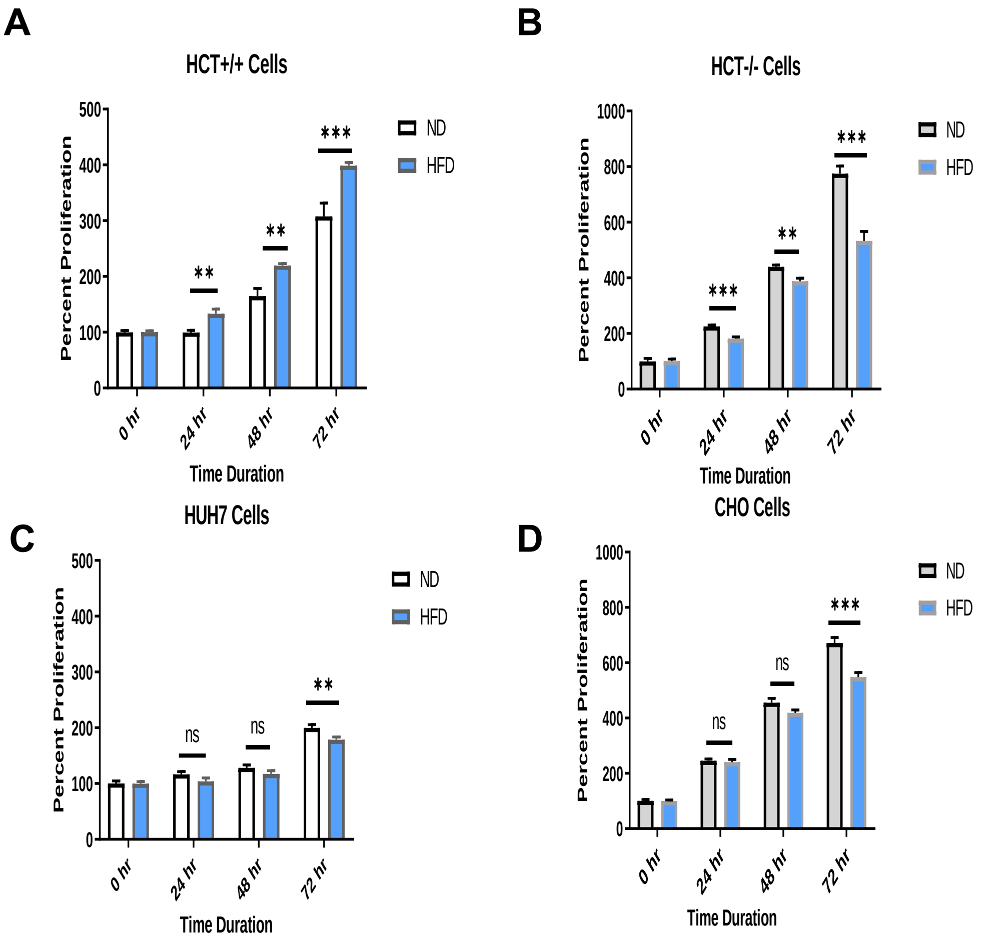


**Figure S4: Analysis of proliferation of cells cultured in ND or HFD serum.**

Cells were cultured in ND and HFD serum-containing media for different time points i.e. 24 hr, 48 hr, 72 hr, and cell growth was analyzed by MTT assay. In brief, cells were seeded at a density of 0.005 × $10^6$ cells per well in 96-well plates and allowed to adhere. After 24 hr, the cells were subjected to 12 hr starvation period. Subsequently, media containing 5% ND or 5% HFD were added to the wells. At different time points, the medium was removed, and MTT was added to wells. The insoluble crystals of MTT were dissolved in isopropanol and absorbance was recorded at 570 nm. The absorbance of cells before replacing the media with ND and HFD serum-containing media was set as 100 percentages and relative to this the graph was plotted. Percent proliferation of cells cultured in ND and HFD serum by MTT assay (A) HCT+/+ (B) of HCT-/- (C) Huh7 (D) CHO. The experiment was performed in triplicates.

**Supplementary figure S5**

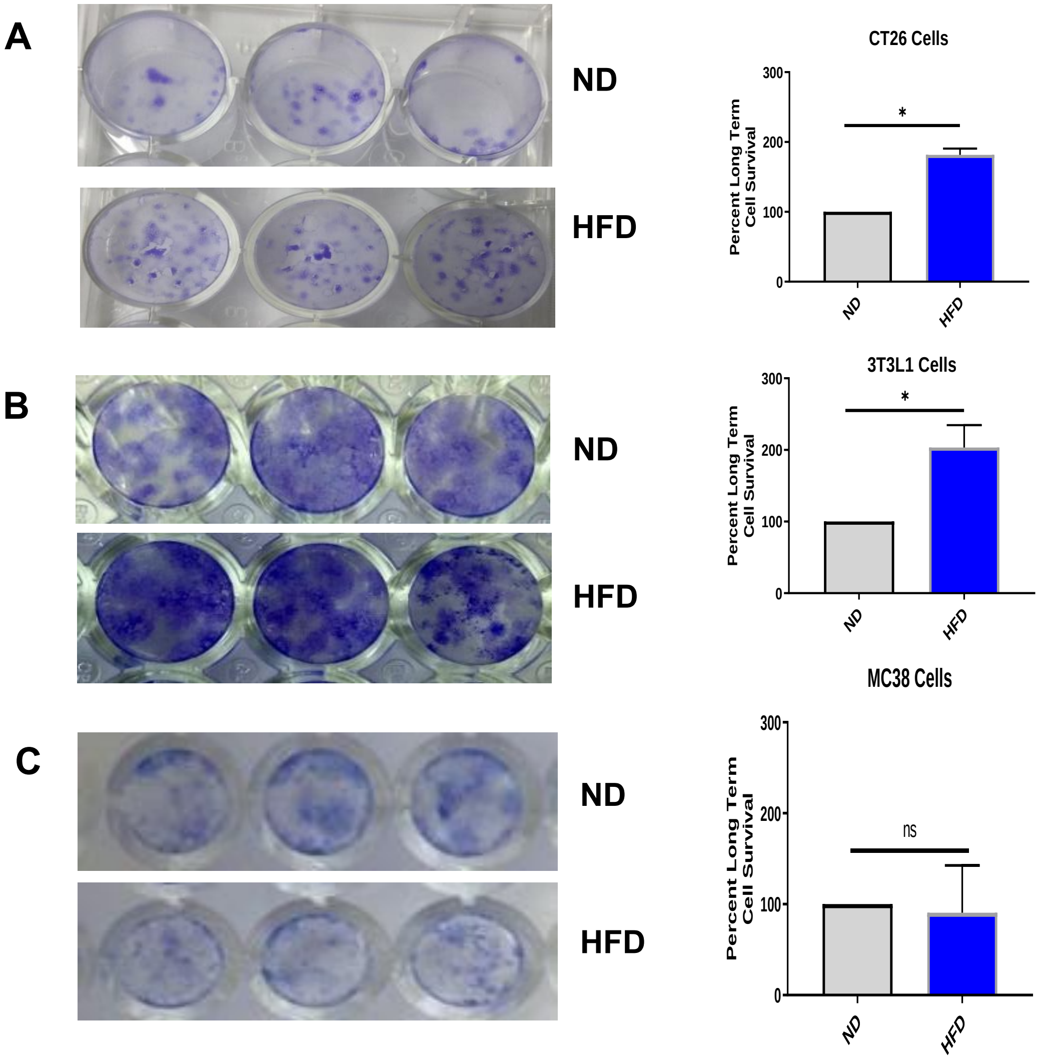

**Figure S5: Analysis of long term growth of cells cultured in ND and HFD serum.**

Cells were cultured in ND and HFD serum-containing media for 7-10 days in ND or HFD serum containing media and allowed to form colonies. The images of crystal violet-stained colonies of cells cultured in ND or HFD serum-containing media along with bar graph showing percent long term survival of cells (A) CT26 (B) 3T3-L1, and (C) MC38. The experiment was performed in triplicates.

**Supplementary figure S6**

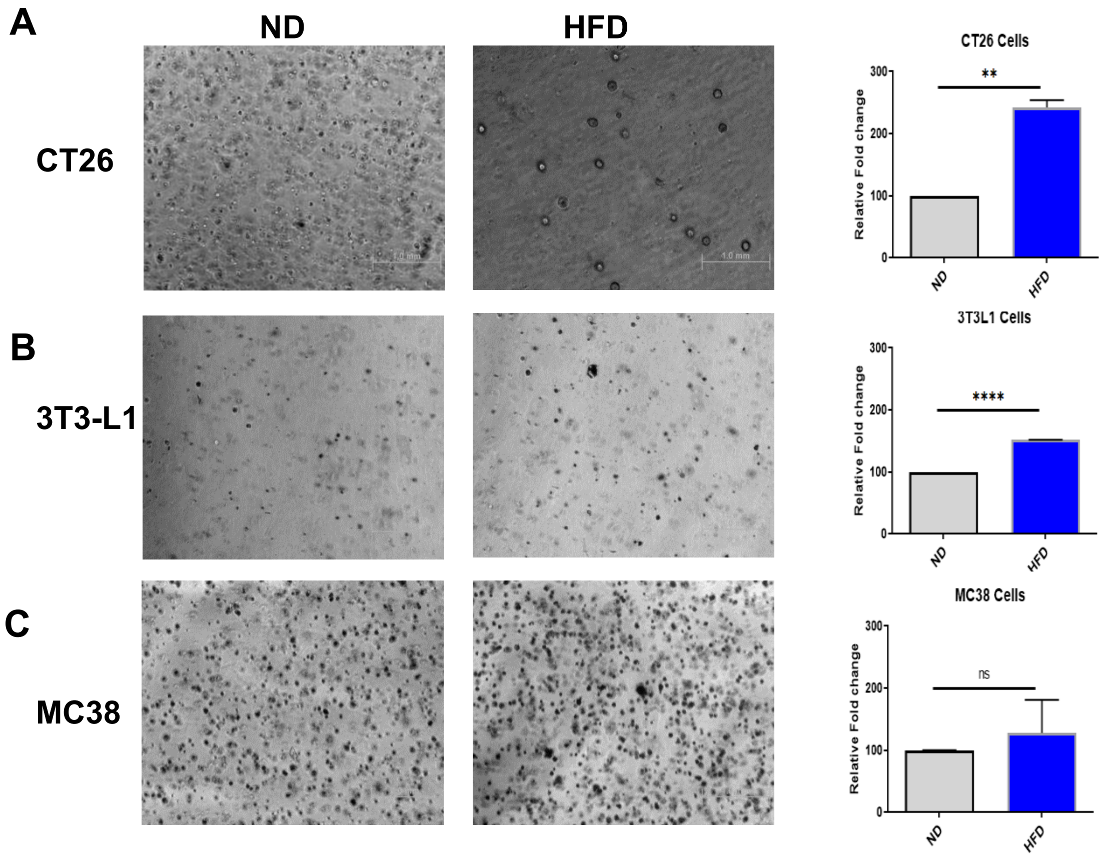


**Figure S6: Analysis of anchorage independent growth of cells in ND and HFD serum-containing media**.

Approximately 1000 cells/well mixed with low melting agarose were layered on normal melting agarose-coated wells. Then 5% ND or 5% HFD serum-containing media was added over the LMA layer and cells were allowed to form colonies for 10-12 days. Phase contrast image of colonies of cells, images were taken at 4X magnification. Quantification of colonies was done through measuring intensity by Image J. (A) CT26 (B) 3T3-L1, and (C) MC38.  The experiment was performed in triplicates and performed twice.

**Supplementary figure S7**

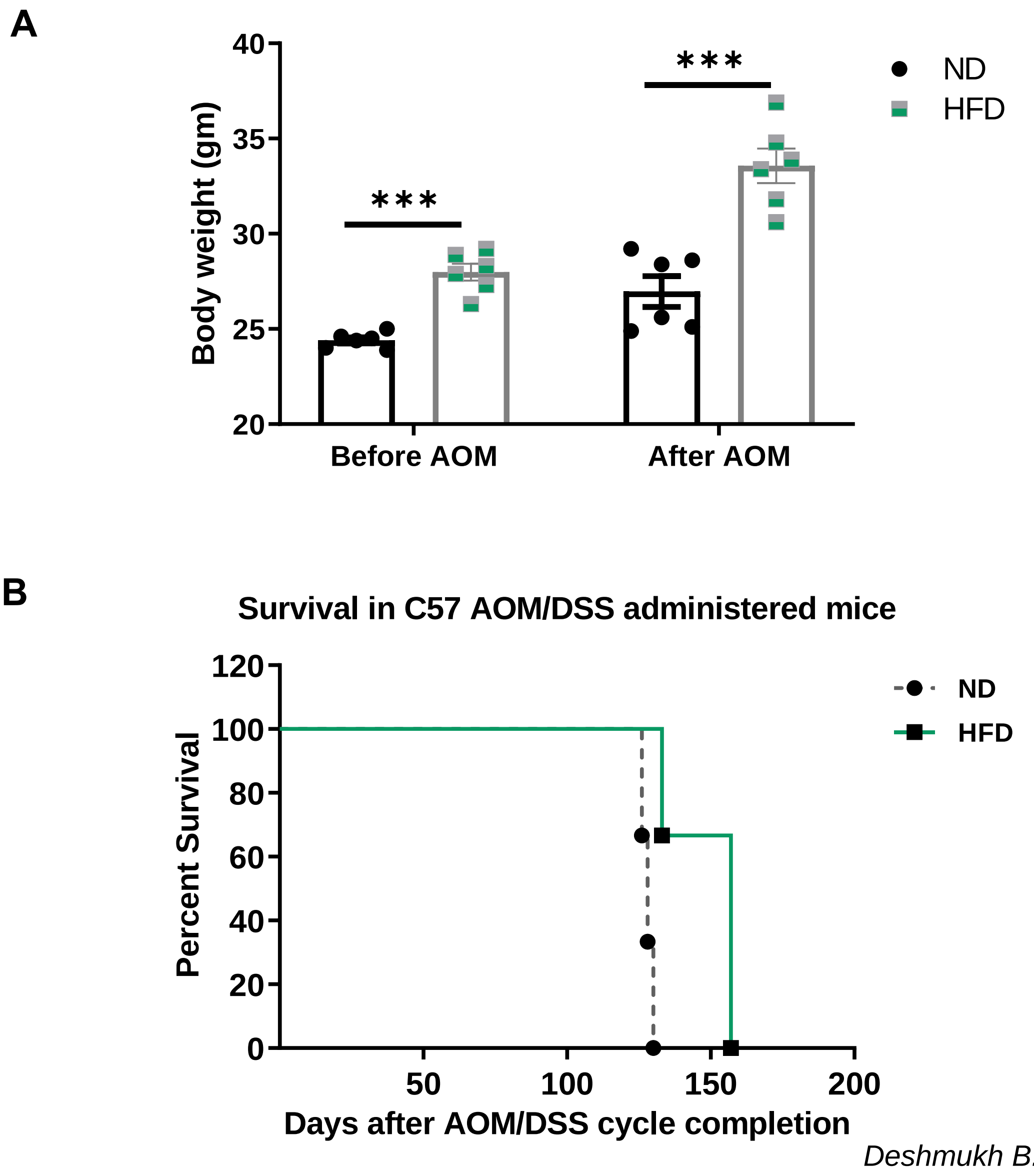

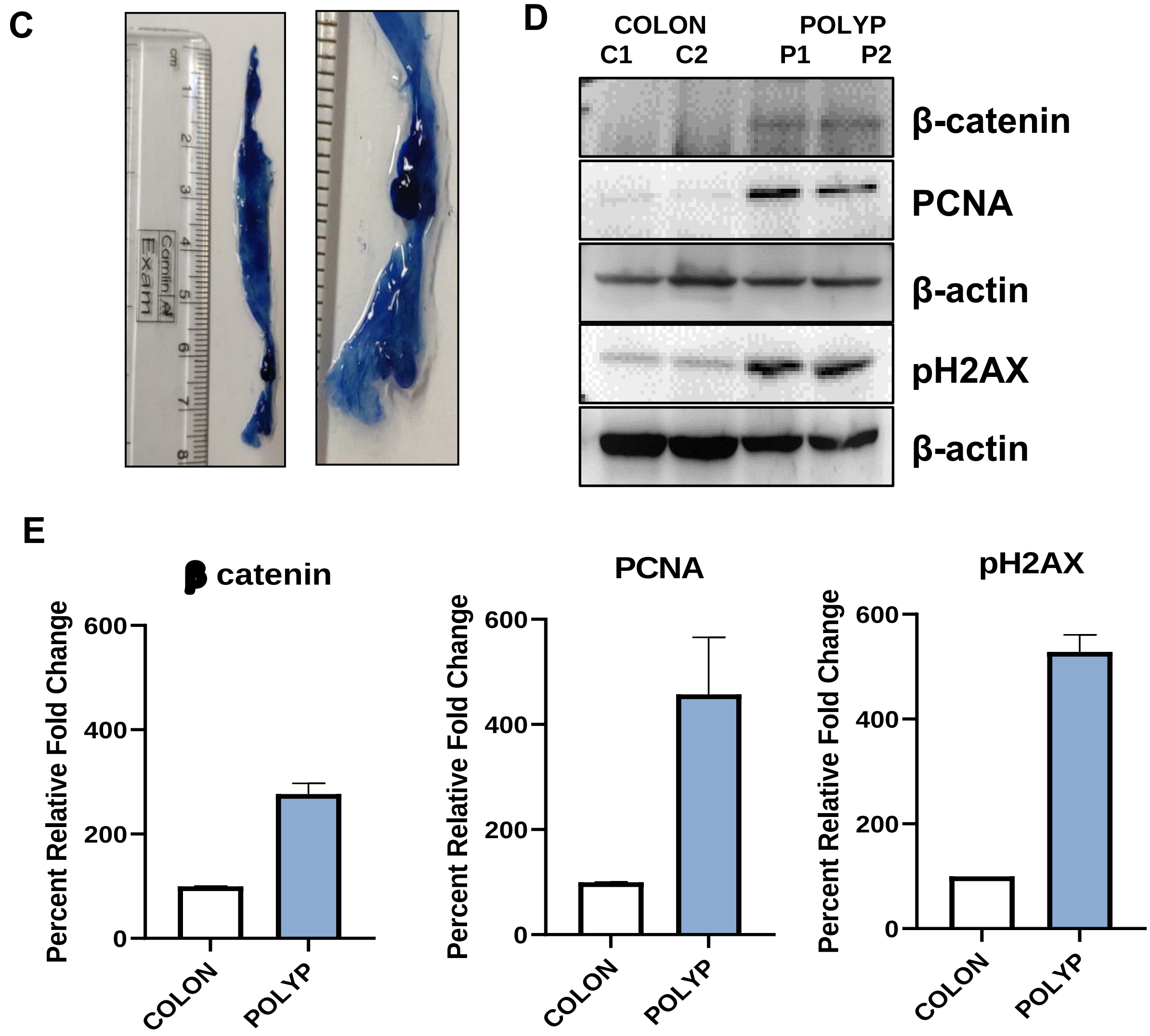


**Figure S7: Effect of AOM administration on weight of mice, survival curve and excised colonic polyps exhibit dysplastic phenotype.**

(A) The weight difference in ND and HFD mice before and after AOM administration. Before AOM administration, the difference between ND and HFD mice was approximately 3.5 gm. This difference was increased at the end of AOM by approximately 7 gm. (B) Survival curve of AOM-administered ND and HFD mice. After completion of the

AOM/DSS cycle, 3 mice from each group were randomly selected for survival analysis which may or may not have polyps in the colon. Two of the HFD mice died on the same day (157 days). The graph shows the percent survival of AOM/DSS administered ND and HFD mice. (C) Dissected colon was fixed with 10% paraformaldehyde and stained with 0.2% methylene blue, which showed the presence of dysplastic polyps. (D) Immuno-blots of β-catenin, PCNA and pH2AX in colon tissue and polyp lysates. (E) The bar graph represents the fold change in protein levels of β-catenin, PCNA and pH2AX.
